# Speech categorization reveals the role of early-stage temporal-coherence processing in auditory scene analysis

**DOI:** 10.1101/2021.09.06.459159

**Authors:** Vibha Viswanathan, Barbara G. Shinn-Cunningham, Michael G. Heinz

## Abstract

Temporal coherence of sound fluctuations across spectral channels is thought to aid auditory grouping and scene segregation. Although prior studies on the neural bases of temporal-coherence processing focused mostly on cortical contributions, neurophysiological evidence suggests that temporal-coherence-based scene analysis may start as early as the cochlear nucleus (i.e., the first auditory region supporting cross-channel processing over a wide frequency range). Accordingly, we hypothesized that aspects of temporal-coherence processing that could be realized in early auditory areas may shape speech understanding in noise. We then explored whether physiologically plausible computational models could account for results from a behavioral experiment that measured consonant categorization in different masking conditions. We tested whether within-channel masking of target-speech modulations predicted consonant confusions across the different conditions, and whether predicted performance was improved by adding across-channel temporal-coherence processing mirroring the computations known to exist in the cochlear nucleus. Consonant confusions provide a rich characterization of error patterns in speech categorization, and are thus crucial for rigorously testing models of speech perception; however, to the best of our knowledge, they have not been utilized in prior studies of scene analysis. We find that within-channel modulation masking can reasonably account for category confusions, but that it fails when temporal fine structure (TFS) cues are unavailable. However, the addition of across-channel temporal-coherence processing significantly improves confusion predictions across all tested conditions. Our results suggest that temporal-coherence processing strongly shapes speech understanding in noise, and that physiological computations that exist early along the auditory pathway may contribute to this process.

## 1 Introduction

An accumulating body of evidence suggests that temporal-coherence processing is important for multisensory scene analysis (Singer and Gray, 1995; Elhilali et al., 2009). In audition, a rich psychophysical literature on grouping (Darwin, 1997), comodulation masking release (CMR; Schooneveldt and Moore, 1987) and cross-channel interference (Apoux and Bacon, 2008), and pitch-based masking release (Oxenham and Simonson, 2009) support the theory that temporally coherent sound modulations can bind together sound elements across distinct spectral channels to form a perceptual object, which can help perceptually separate different sources in an acoustic mixture. This theory may help explain how we perform speech separation in a multi-source environment (Krishnan et al., 2014), as speech naturally has common temporal fluctuations across different channels, particularly in the syllabic (0–5 Hz), phonemic (5–64 Hz), and periodicity (i.e., pitch; 64–300 Hz) ranges (Crouzet and Ainsworth, 2001; Swaminathan and Heinz, 2011).

Prior studies on the neural bases of temporal-coherence processing mostly focused on cortical contributions (Elhilali et al., 2009; Teki et al., 2013; O’Sullivan et al., 2015). However, single-unit measurements and computational modeling of across-channel CMR effects suggest that temporal-coherence-based scene analysis may start early in the auditory pathway; for instance, the cochlear nucleus has the physiological mechanisms (e.g., wideband inhibition) needed to support such analysis (Pressnitzer et al., 2001; Meddis et al., 2002). Moreover, attention, which operates on segregated auditory objects (Shinn-Cunningham, 2008), affects responses in the primary auditory cortex (Hillyard et al., 1973). Given this, binding and scene segregation likely start even earlier, such as brainstem, and accumulate along the auditory pathway. However, no prior studies have directly tested the theory that speech understanding in noise may be shaped by aspects of temporal-coherence processing that exist in early auditory areas.

While previous studies of temporal-coherence processing mostly used non-speech stimuli (e.g., Elhilali et al., 2009; Teki et al., 2013; O’Sullivan et al., 2015), a parallel literature on modeling speech-intelligibility mechanisms typically focused on overall intelligibility to test predictions of performance (Jørgensen et al., 2013; Relaño-Iborra et al., 2016). A detailed characterization of error patterns in speech categorization—crucial in order to rigorously examine any theory of speech perception—has not been previously used in studies of scene analysis. In contrast, confusion patterns in speech categorization, such as consonant/vowel confusion matrices (Miller and Nicely, 1955), have been widely used in the speech acoustics and cue-weighting literatures, and can indeed provide deeper insight into underlying mechanisms if utilized to test theories of scene analysis.

To address these gaps, we used a combination of online consonant-identification experiments and computational modeling of temporal-coherence processing that is physiologically plausible in the cochlear nucleus (Pressnitzer et al., 2001), the first auditory area where cross-channel processing over a wide frequency range is supported. We asked whether the masking of target-speech envelopes by distracting masker modulations (i.e., modulation masking; Bacon and Grantham, 1989; Stone and Moore, 2014) within individual frequency channels (as implemented in current speech-intelligibility models; Jørgensen et al., 2013; Relaño-Iborra et al., 2016) is sufficient to predict consonant categorization, or if across-channel temporal-coherence processing improves predictions by accounting for interference from masker elements that are temporally coherent with target elements but in different frequency channels. Crucially, instead of just trying to predict perceptual intelligibility measurements from model outputs, we predicted consonant confusion patterns in various listening conditions. Considering the error patterns in consonant categorization (i.e., when an error was made, what consonant was reported instead of the consonant presented) provided a richer characterization of the processes engaged during speech perception compared to looking only at percent-correct scores. Our combined use of consonant confusions and physiologically plausible computational modeling provides independent evidence for the role of temporal-coherence processing in scene analysis and speech perception. Moreover, it suggests that this processing may start earlier in the auditory pathway than previously thought.

## 2 Materials and Methods

### 2.1 Stimulus generation

The stimuli used in the present study draw from and expand on the Materials and Methods previously described in Viswanathan et al. (2021b). 20 consonants from the STeVI corpus (Sensimetrics Corporation, Malden, MA) were used. The consonants were /b/, 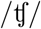, /d/, /ð/, /f/, /g/, 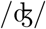, /k/, /l/, /m/, /n/, /p/, /r/, /s/, /∫/, /t/, /θ/, /v/, /z/, and /3/. The consonants were presented in CV (consonant-vowel) context, where the vowel was always /a/. Each consonant was spoken by two female and two male talkers (to reflect real-life talker variability). The CV utterances were embedded in the carrier phrase: “You will mark /CV/ please” (i.e., in natural running speech). Stimuli were created for five experimental conditions:

1. **Speech in Quiet (SiQuiet):** Speech in quiet was used as a control condition.
2. **Speech in Speech-shaped Stationary Noise (SiSSN):** Speech was added to stationary Gaussian noise at -8 dB signal-to-noise ratio (SNR). The long-term spectra of the target speech (including the carrier phrase) and that of stationary noise were adjusted to match the average (across instances) long-term spectrum of the four-talker babble. A different realization of stationary noise was used for each SiSSN stimulus.
3. **Speech in Babble (SiB):** Speech was added to four-talker babble at -8 dB SNR. The long-term spectrum of the target speech (including the carrier phrase) was adjusted to match the average (across instances) long-term spectrum of the four-talker babble. Each SiB stimulus was created by randomly selecting a babble sample from a list comprising 72 different four-talker babble maskers obtained from the QuickSIN corpus (Killion et al., 2004).
4. **Speech in a masker with only DC modulations (SiDCmod) (Stone et al., 2012):** In line with the procedure described in Stone et al. (2012), the target speech was filtered into 28 channels between 100–7800 Hz and a sinusoidal masker centered on each channel was added to the channel signal at -18 dB SNR. To minimize peripheral interactions between maskers, odd-numbered channels were presented to one ear and even to the other; this procedure effectively yields an unmodulated masker (i.e., a masker with a modulation spectrum containing only a DC component). Thus, the SiDCmod condition presented stimuli that were dichotic, unlike the other conditions, which presented diotic stimuli. The long-term spectra of the target speech (including the carrier phrase) and that of the masker were adjusted to match the average (across instances) long-term spectrum of the four-talker babble.
5. **Vocoded Speech in Babble (Vocoded SiB):** SiB at 0 dB SNR was subjected to 64-channel envelope vocoding. A randomly selected babble sample was used for each Vocoded SiB stimulus, similar to what was done for intact SiB. In accordance with the procedure described in Viswanathan et al. (2021b), we retained the cochlear-level envelopes during vocoding but replaced the stimulus temporal fine structure (TFS) with a noise carrier. We verified that the vocoding procedure did not significantly change envelopes at the cochlear level, as described in Viswanathan et al. (2021b).

Table 1 describes the rationale behind including these different stimulus conditions in our study.

**Table 1:** Rationale for the different stimulus conditions included in this study. The different listening conditions were chosen to span a range of modulation masking spectral profiles and temporal fine structure (TFS) information, which allows for theories of scene analysis based on within-channel modulation masking and across-channel temporal coherence to be tested in a rigorous manner. Collectively these conditions represent a diversity of scene acoustics, including important examples in our environment and clinical applications. The SNR levels were chosen to give approximately equal overall intelligibility across SiSSN, SiB, SiDCmod, and Vocoded SiB using a behavioral pilot study with three subjects who did not participate in the online consonant identification experiment. This was done to obtain roughly equal variance in the consonant confusion estimates for these conditions, which allows us to fairly compare confusion patterns across them. Equalizing intelligibility also maximizes the statistical power for detecting differences in the pattern of confusions. The overall intelligibility in each of these conditions was ~60%, which yielded a sufficient number of confusions for analysis.

| No. | Stimulus condition | Rationale for inclusion in study |
| --- | --- | --- |
| 1 | Speech in Quiet (SiQuiet) | Used as a control condition |
| 2 | Speech in Speech-shaped Stationary Noise (SiSSN) at -8 dB SNR | Widely used in the literature; used for calibration of prediction model |
| 3 | Speech in Babble (SiB) at -8 dB SNR | Simulates ecologically relevant cocktail-party listening |
| 4 | Speech in a masker with only DC modulations (SiDCmod) at -18 dB SNR | To obtain a different modulation masking profile from stationary noise (which contains relatively more high-frequency modulation energy) and babble (which contains relatively more low-frequency modulation power) (Viswanathan et al., 2021a) |
| 5 | SiB at 0 dB SNR subjected to 64-channel envelope vocoding | Used to compare performance across models that consider TFS and those that do not (since TFS can influence scene analysis and can convey consonant voicing information in noise; Viswanathan et al., 2021a,b) |

The stimulus used for online volume adjustment was running speech mixed with four-talker babble. The speech and babble samples were obtained from the QuickSIN corpus (Killion et al., 2004); these were repeated over time to obtain a *∼*20 s total stimulus duration (to give subjects sufficient time to adjust their computer volume with the instructions described in Section 2.3). The root mean square (RMS) value of this stimulus corresponded to 75% of the dB difference between the softest and loudest stimuli in the consonant identification experiment, which ensured that no stimulus was too loud for subjects once they had adjusted their computer volume to a comfortable level.

### 2.2 Participants

Full details of participant recruitment and screening are provided in Viswanathan et al. (2021b), and are only briefly reviewed here. Anonymous subjects were recruited for online data collection using Prolific.co. A three-part subject-screening protocol developed and validated by Mok et al. (2021) was used to restrict the subject pool. This protocol included a survey on age, native-speaker status, presence of persistent tinnitus, and history of hearing and neurological diagnoses, followed by headphone/earphone checks and a speech-in-babble-based hearing screening. Subjects who passed this screening protocol were invited to participate in the consonant identification study, and when they returned, headphone/earphone checks were performed again. Only subjects who satisfied the following criteria passed the screening protocol: (i) 18–55 years old, (ii) self-reported no hearing loss, neurological disorders, or persistent tinnitus, (iii) born and residing in US/Canada, and native speaker of North American English, (iv) experienced Prolific subject, and (v) passed the headphone/earphone checks and speech-in-babble-based hearing screening (Mok et al., 2021). Subjects provided informed consent in accordance with remote testing protocols approved by the Purdue University Institutional Review Board (IRB).

### 2.3 Experimental design

The online consonant identification experiment was previously described in Viswanathan et al. (2021b). Subjects performed the experiment using their personal computers and headphones/earphones. Our online infrastructure included checks to prevent the use of mobile devices. The experiment had three parts: (i) Headphone/earphone checks, (ii) Demonstration (“Demo”), and (iii) Test. Each of these three parts had a volume-adjustment task at the beginning. In this task, subjects were asked to make sure that they were in a quiet room and wearing wired (not wireless) headphones or earphones. They were instructed not to use desktop/laptop speakers. Headphone use was checked using the procedures outlined in Mok et al. (2021). They were then asked to set their computer volume to 10–20% of the full volume, following which they were played a speech-in-babble stimulus and asked to adjust their volume up to a comfortable but not too loud level. Once subjects had adjusted their computer volume, they were instructed not to adjust the volume during the experiment, as that could lead to sounds being too loud or soft.

The Demo stage consisted of a short training task designed to familiarize subjects with how each consonant sounds and with the consonant-identification paradigm. Subjects were instructed that in each trial they would hear a voice say “You will mark *something* please.” They were told that they would be given a set of options for *something* at the end of the trial, and that they should click on the corresponding option. After subjects had heard all consonants sequentially (i.e., the same order as the response choices) in quiet, they were tasked with identifying consonants presented in random order and spanning the same set of listening conditions as the Test stage. Subjects were instructed to ignore any background noise and only listen to the particular voice saying “You will mark *something* please.” In order to ensure that all subjects understood and were able to perform the task, only those subjects who scored *≥* 85% in the Demo’s Speech in Quiet control condition were selected for the Test stage.

Subjects were given similar instructions in the Test stage as in the Demo, but told to expect trials with background noise from the beginning. The Test stage presented, in random order, the 20 consonants (with one stimulus repetition per consonant) across all four talkers and all five experimental conditions. In both Demo and Test, the masking noise, when present, started 1 s before the target speech and continued for the entire duration of the trial. This was done to cue the subjects’ attention to the stimulus before the target sentence was played. In both the Demo and Test parts, subjects received feedback after every trial as to whether or not their response was correct to promote engagement with the task. However, subjects were not told what consonant was presented to avoid over-training to the acoustics of how each consonant sounded across the different conditions; the only exception to this rule was in the first sub-part of the Demo where subjects heard all consonants in quiet in sequential order.

We used 50 subjects per talker (subject overlap between talkers was not controlled); with four talkers, this yielded 200 subject-talker pairs, or samples. Separate studies were posted on Prolific.co for the different talkers; thus, when a subject performed a particular study, they would be presented with the speech stimuli for one specific talker consistently over all trials. Within each talker and condition, all subjects performed the task with the same stimuli. Moreover, all condition effect contrasts were computed on a within-subject basis, and averaged across subjects.

### 2.4 Data preprocessing

Only samples with intelligibility scores *≥* 85% for the Speech in Quiet control condition in the Test stage were included in results reported here. All conditions for the remaining samples were excluded from further analyses as a data quality control measure. This yielded a final N=191 samples.

### 2.5 Quantifying confusion matrices from perceptual measurements

The 20 English consonants used in this study were assigned the phonetic features described in Table 2. The identification data collected in the Test stage were used to construct consonant confusion matrices (pooled over samples) for the different conditions; these matrices in turn were used to construct voicing, place of articulation (POA), and manner of articulation (MOA) confusion matrices by pooling over all consonants.

**Table 2:**
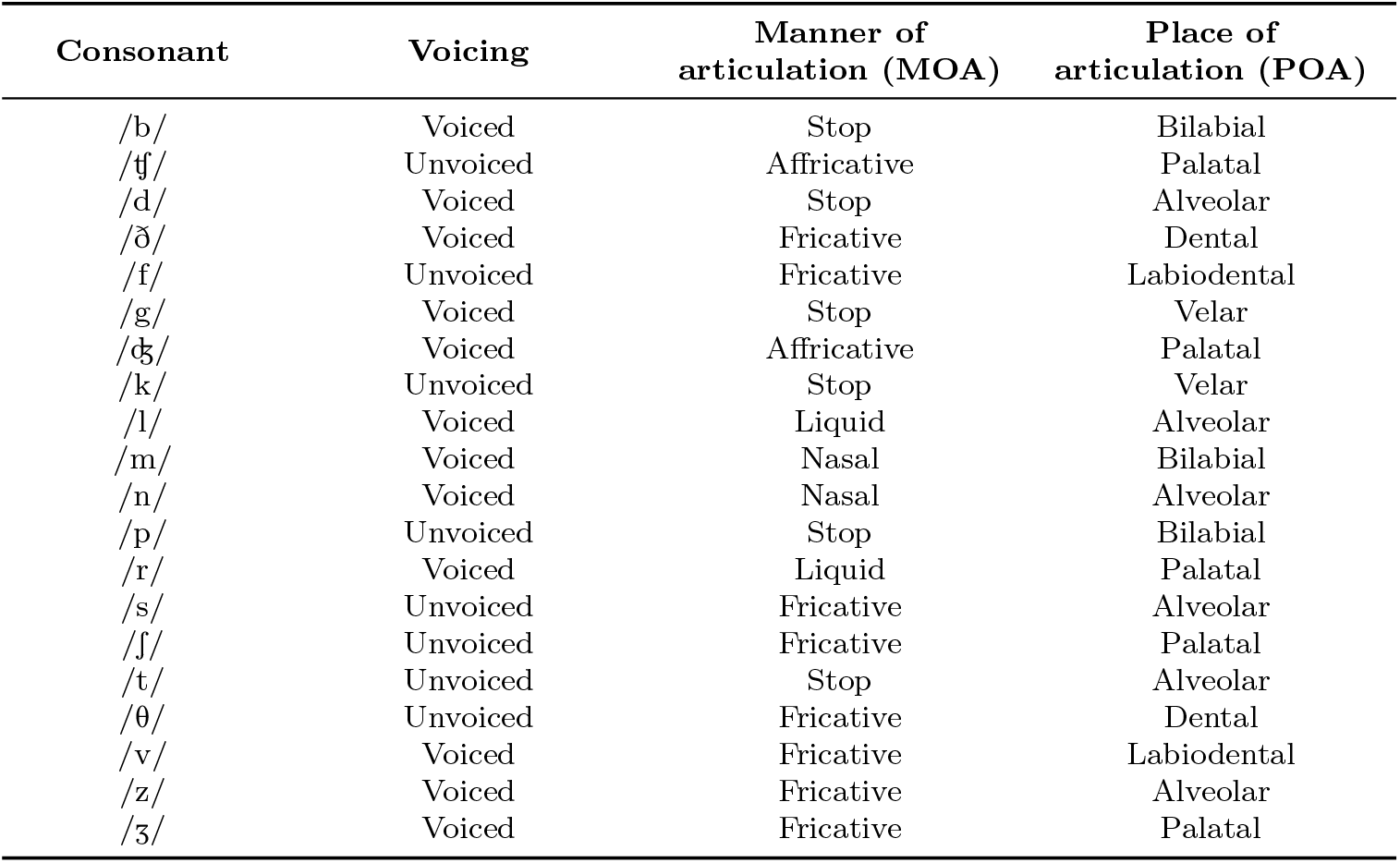
Phonetic features of the 20 English consonants used in this study.

In order to test whether there are significant differences in confusion patterns between SiSSN, SiB, SiDCmod, and Vocoded SiB, we first normalized the overall intelligibility for these conditions to 60% by scaling the consonant confusion matrices such that the sum of the diagonal entries was the desired intelligibility (note that overall intelligibility was not normalized for the main modeling analyses of this study). By matching intelligibility in this manner, differences in confusion matrices across conditions could be attributed to changes in consonant categorization and category errors rather than differences in overall error counts (due to one condition being inherently easier at a particular SNR). Since overall intelligibility was similar across conditions to start with (Fig. 5), small condition differences in intelligibility could be normalized without loss of statistical power. Confusion-matrix differences between the intelligibility-matched conditions were then compared with appropriate null distributions of zero differences (see Section 2.8) to extract statistically significant differences (shown in Fig. 6).

### 2.6 Auditory periphery modeling

The auditory-nerve model of Bruce et al. (2018) was used to simulate processing by the auditory periphery. The parameters of this model were set as follows. 30 cochlear filters with characteristic frequencies (CFs) equally spaced on an ERB-number scale (Glasberg and Moore, 1990) between 125 Hz and 8 kHz were used. Normal function was chosen for the outer and inner hair cells. The species was chosen to be human with the Shera et al. (2002) cochlear tuning at low sound levels; however, with suppression, the Glasberg and Moore (1990) tuning is effectively obtained for our broad-band, moderate-level stimuli (Heinz et al., 2002; Oxenham and Shera, 2003). The noise type parameter for the inner-hair-cell synapse model was set to fixed fractional Gaussian noise to yield a constant spontaneous auditory-nerve firing rate. To avoid single-fiber saturation effects, the spontaneous rate of the auditory-nerve fiber was set to 10, corresponding to that of a medium-spontaneous-rate fiber. An approximate implementation of the power-law adaptation dynamics in the synapse was used. The absolute and relative refractory periods were set to 0.6 ms.

The periphery model was simulated with the same speech stimuli used in our psychophysical experiment (i.e., CV utterances that spanned 20 consonants, four talkers, and five conditions, and were embedded in a carrier phrase) as input. The level for the target speech was set to 60 dB SPL across all stimuli, as this produced sufficient (i.e., firing rate greater than spontaneous rate) model auditory-nerve responses for consonants in quiet and also did not saturate the response to the loudest stimulus. The periphery model was provided with just one audio channel input for all conditions except SiDCmod, as that was the only condition that was dichotic rather than diotic. Instead, for SiDCmod, the model was separately simulated for each of the two audio channels. 200 stimulus repetitions were used to derive peri-stimulus time histograms (PSTHs) from model auditory-nerve outputs. The model was simulated for the full duration of each stimulus (versus just the time period when the target consonant was presented). A PSTH bin width of 1 ms (i.e., a sampling rate of 1 kHz) was used. This was done so as to capture fine-structure phase locking up to and including the typical frequency range of human pitch for voiced sounds. In the case of the SiDCmod condition, a separate PSTH was computed for each of the two dichotic audio channels.

Although the full speech stimuli (including the carrier phrase and CV utterances) were used as inputs to the periphery model, the responses to the target consonants were manually segmented out from the model PSTHs before being input into the scene analysis models. To do this, we calculated the time segment corresponding to when the target consonant was presented for each speech-in-quiet stimulus by visualizing speech spectrograms computed by gammatone filtering (Patterson et al., 1987) followed by Hilbert-envelope extraction (Hilbert, 1906). 128 gammatone filters were used for this purpose, with center frequencies between 100–8000 Hz and equally spaced on an ERB-number scale (Glasberg and Moore, 1990). A fixed duration of 104.2 ms was used for each consonant segment. Segmentation accuracy was verified by listening to the segmented consonant utterances. The time segments thus derived were used to extract model auditory-nerve responses to the different target consonants across the different conditions and talkers. These responses were then used as inputs to the scene analysis models described below.

### 2.7 Scene analysis modeling to predict consonant confusions

In order to study the contribution of across-channel temporal-coherence processing to consonant categorization, we constructed two different scene analysis models. The first is a within-channel modulation-masking-based scene analysis model inspired by Relaño-Iborra et al. (2016), and the second is a simple across-channel temporal coherence model mirroring the physiological computations that are known to exist in the cochlear nucleus (Pressnitzer et al., 2001).

In the within-channel modulation-masking-based model, the auditory-nerve PSTHs (i.e., the outputs from the periphery model; see Section 2.6) corresponding to the different consonants, conditions, and talkers were filtered within a 1-ERB bandwidth (Glasberg and Moore, 1990) to extract band-specific envelopes. Note that the envelopes extracted from auditory-nerve outputs may contain some TFS converted to envelopes via inner-hair-cell rectification (assuming envelope and TFS are defined at the output of the cochlea), but that is the processing that is naturally performed by the auditory system as well. Pairwise dynamic time warping (Rabiner, 1993) was performed to align the results for each pair of consonants across time. Dynamic time warping can help compensate for variations in speaking rate across consonants. A modulation filterbank (Ewert and Dau, 2000; Jørgensen et al., 2013) was then used to decompose the results at each CF into different modulation frequency (MF) bands. This filterbank consists of a low-pass filter with a cutoff frequency of 1 Hz in parallel with eight band-pass filters with octave spacing, a quality factor of 1, and center frequencies ranging from 2 to 256 Hz. For each condition, talker, CF, MF, and consonant, Pearson correlation coefficients were computed between the filterbank output for that consonant in that particular condition and the output for each of all 20 consonants in quiet. Each of the individual correlations was squared to obtain the variance explained; the results were averaged across talkers, CFs, and MFs to obtain a “raw” neural metric *ψ* for each experimental condition. A separate *ψ* value was obtained for each condition, and every pair of consonant presented and option for consonant reported. For the dichotic SiDCmod condition, the variance explained was separately computed for the left and right ears at each CF, then the maximum across the two ears (i.e., the “better-ear” contribution) was used for that CF (Zurek, 1993). Finally, for each condition, the *ψ* values were normalized such that their sum across all options for consonants reported for a particular consonant presented was equal to 1; this procedure yielded a condition-specific “neural consonant confusion matrix.”

We wanted to test whether across-channel temporal-coherence processing of input fluctuations could better predict consonant categorization than a purely within-channel modulation masking model. To simulate across-channel temporal-coherence processing, we modeled a physiologically plausible wideband-inhibition-based temporal-coherence processing circuit proposed by Pressnitzer et al. (2001) to account for physiological correlates of CMR in the cochlear nucleus. A schematic of this circuit is provided in Figure 1. The overall across-channel scene analysis model is similar to the within-channel model, except that the envelope extraction stage of the within-channel model is replaced with the CMR circuit model in the across-channel model. Thus, the across-channel model can account for both within-channel modulation masking effects as well as across-channel temporal-coherence processing. Figure 2 shows schematics of both the within- and across-channel models.

**Figure 1:**
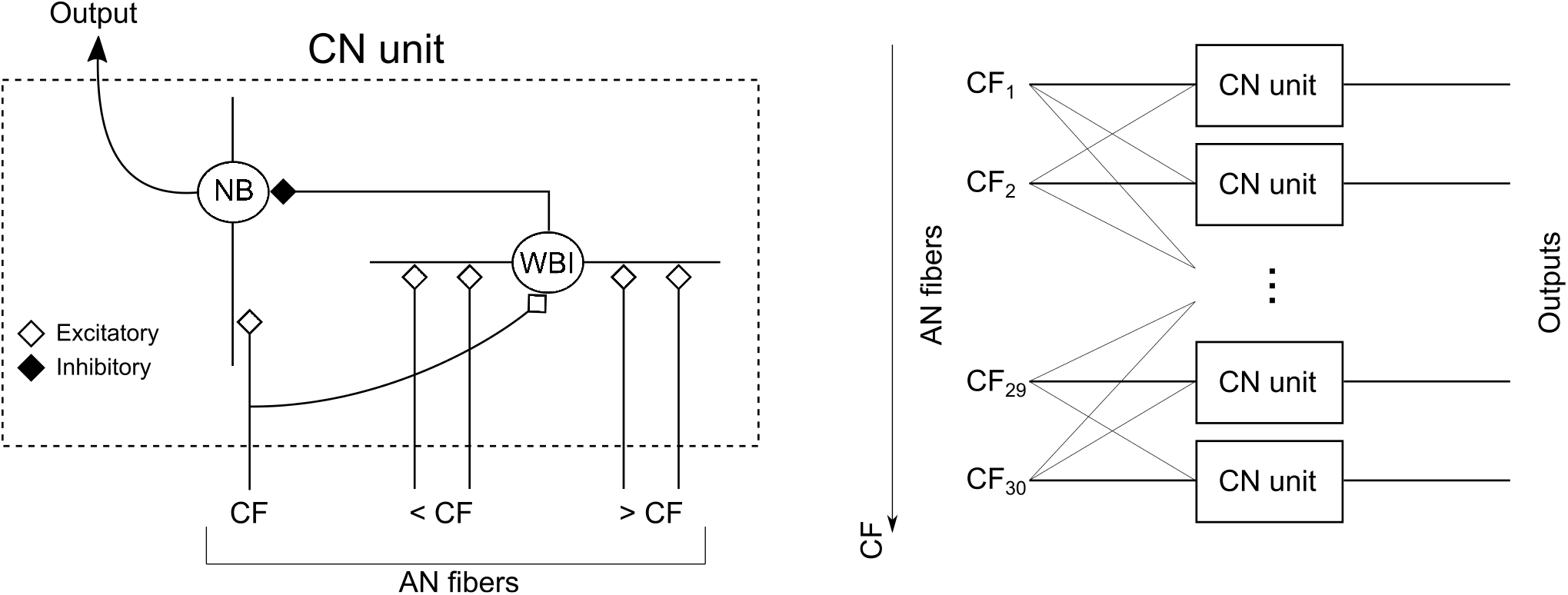
Comodulation masking release (CMR) circuit based on wideband inhibition in the cochlear nucleus. This physiologically plausible circuit was proposed by Pressnitzer et al. (2001) to model CMR effects seen in the cochlear nucleus (CN). CN units at different characteristic frequencies (CFs) form the building blocks of this circuit. Each CN unit consists of a narrowband cell (NB) that receives narrow on-CF excitatory input from the auditory nerve (AN) and inhibitory input from a wideband inhibitor (WBI). The WBI in turn receives excitatory inputs from AN fibers tuned to CFs spanning two octaves below to one octave above the CF of the NB that it inhibits. The time constants for the excitatory and inhibitory synapses are 5 ms and 1 ms, respectively. The WBI input to the NB is delayed with respect to the AN input by 2 ms. Note that our model simulations were rate-based, i.e., they used AN peri-stimulus time histograms (PSTHs) rather than spikes. Thus, all outputs were half-wave rectified (i.e., firing rates were positive at every stage). All synaptic filters were initially normalized to have unit gain, then the gain of the inhibitory input was allowed to vary parametrically to implement different excitation-to-inhibition (EI) ratios between 3:1 and 1:1. The EI ratio was adjusted so as to obtain the best consonant confusion prediction accuracy for SiSSN (i.e., the calibration condition), and the optimal ratio for the calibration condition was found to be 1.75:1. Note that the model parameter corresponding to the EI ratio cannot be readily compared to its physiological correlate because the model is rate-based and lacks important membrane conductance properties that spiking models can be endowed with.

**Figure 2:**
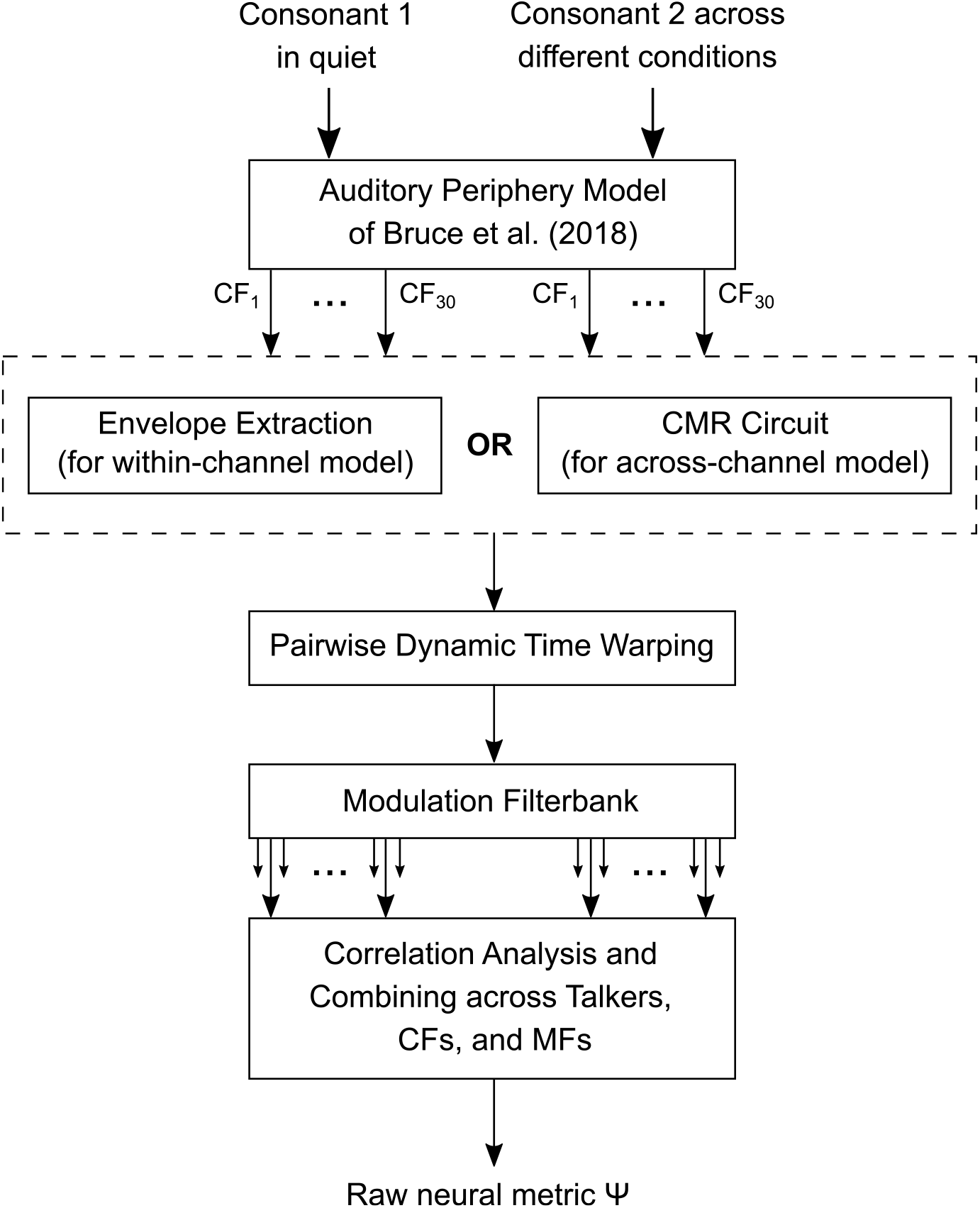
Schematic of the within- and across-channel scene analysis models. The speech stimuli were input into the Bruce et al. (2018) model, which simulated a normal auditory periphery with 30 cochlear filters having characteristic frequencies (CFs) equally spaced on an ERB-number scale (Glasberg and Moore, 1990) between 125 Hz and 8 kHz. PSTHs from the periphery model were processed to retain only the time segments when the target consonants were presented. For the within-channel model, these results were filtered within a 1-ERB bandwidth (Glasberg and Moore, 1990) to extract band-specific envelopes; however, for the across-channel model, the results were instead input to the CMR circuit model (Fig. 1). Pairwise dynamic time warping was performed to align the outputs from the previous step across time for each pair of consonants. A modulation filterbank (Ewert and Dau, 2000; Jørgensen et al., 2013) was then used to decompose the results at each CF into different modulation frequency (MF) bands. This filterbank consists of a low-pass filter with a 1-Hz cutoff in parallel with eight band-pass filters with octave spacing, a quality factor of 1, and center frequencies between 2–256 Hz. For each condition, talker, CF, MF, and consonant, Pearson correlation coefficients were computed between the filterbank output for that consonant in that particular condition and the output for each of all 20 consonants in quiet. Each of the individual correlations was squared to obtain the variance explained; the results were averaged across talkers, CFs, and MFs to obtain a “raw” neural metric *ψ* for each experimental condition. A separate *ψ* value was obtained for each condition, and every pair of consonant presented and option for consonant reported. The *ψ* values were normalized such that their sum across all options for consonants reported for a particular consonant presented was equal to 1, which yielded a condition-specific “neural consonant confusion matrix.”

To verify that the CMR circuit model (Fig. 1) produced physiological correlates of CMR similar to those reported by Pressnitzer et al. (2001), we used the same complex stimuli that they used (Fig. 3). The stimuli consisted of a target signal in a 100% sinusoidally amplitude-modulated (SAM) tonal complex masker. There were three experimental conditions: Reference, Comodulated, and Codeviant. In the Reference condition, the masker had just one component: a SAM tone with a carrier frequency of 1.1 kHz (to allow comparison to data from Pressnitzer et al., 2001); this masking component is also referred to as the on-frequency component (OFC). The Comodulated and Codeviant conditions presented the OFC along with six flanking components that were SAM tones at the same level as the OFC. The carrier frequency separation between the different flanking components and the OFC were -800 Hz, -600 Hz, -400 Hz, 400 Hz, 600 Hz, and 800 Hz, respectively. The flanking components were modulated in phase with the OFC in the Comodulated condition, and 180° out of phase with the OFC in the Codeviant condition. A 10 Hz modulation rate was used for all SAM tones. The target signal consisted of a 50-ms-long (i.e., half of the modulation time period) tone pip at 1.1 kHz that was presented in the dips of the OFC modulation during the last 0.3 s of the stimulus period (i.e., in the last three dips) at different values of signal-to-component ratio (SCR; defined as the signal maximum amplitude over the amplitude of the OFC before modulation). These stimuli were presented to the periphery model, and the corresponding model outputs were passed into the CMR circuit model.

**Figure 3:**
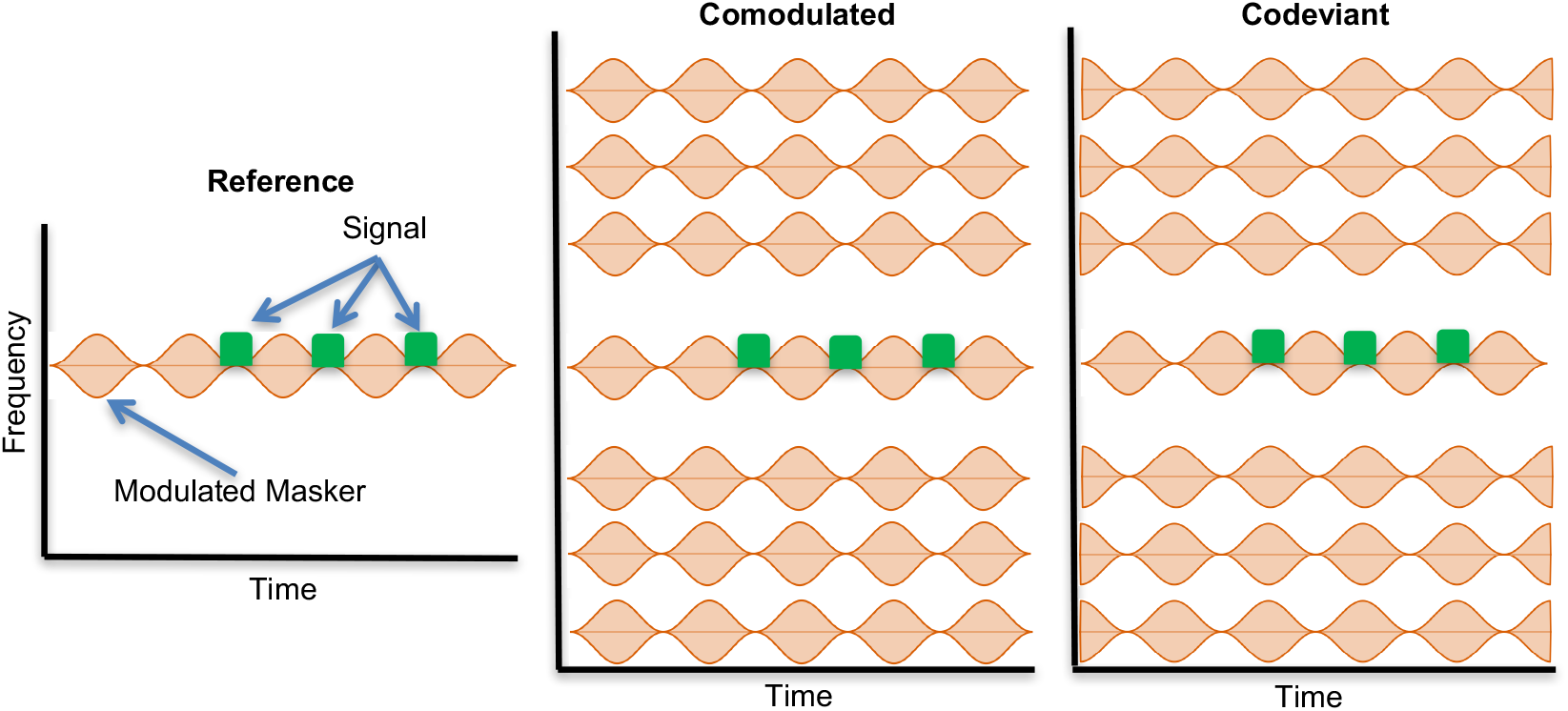
Stimuli used to validate the CMR circuit model. The stimuli used were from Pressnitzer et al. (2001), and consisted of a target signal in a 10-Hz 100% sinusoidally amplitude-modulated (SAM) tonal complex masker. The masker differed depending on the experimental condition. In the Reference condition, the masker was a 1.1 kHz-carrier SAM tone (referred to as the on-frequency component or OFC). In the Comodulated and Codeviant conditions, six flanking components were presented in addition to the OFC. The flanking components were SAM tones at the same level as the OFC. The flanking components were separated from the OFC by -800 Hz, -600 Hz, -400 Hz, 400 Hz, 600 Hz, and 800 Hz, respectively. The modulation of each flanking component was in phase with the OFC modulation in the Comodulated condition, but 180° out of phase with the OFC modulation in the Codeviant condition. The target signal was a 50-ms-long 1.1 kHz tone pip that was presented in the dips of the OFC modulation during the last 0.3 s of the stimulus period (i.e., in the last three dips) at different values of signal-to-component ratio (SCR; defined as the signal maximum amplitude over the amplitude of the OFC before modulation).

The rate-level function at the output of the CMR circuit model (Fig. 4D) closely matches physiological data for chopper units in the ventral cochlear nucleus (Winter and Palmer, 1990), and was used to set the masker level for the CMR stimuli. The firing-rate threshold was 0 dB SPL for pure-tone inputs at CF; thus, a fixed level of 40 dB SPL (i.e., 40 dB SL) was used for the OFC. The PSTH outputs from the CMR circuit model (at 1.1 kHz CF) are shown in Figure 4A. The time-averaged statistics of the firing rate during the last 0.3 s of the stimulus period and in the absence of the target signal were used as the null distribution against which the neurometric sensitivity, *d*′, was calculated; a separate null distribution was derived for each condition. The average firing rate during the target signal periods was compared to the corresponding null distribution to estimate a separate *d*′ for each SCR and condition (Fig. 4B). *d*′ of 0.4 was used to calculate SCR thresholds and the corresponding CMR (threshold difference between the Codeviant and Comodulated conditions). Note that the absolute *d*′ values cannot be interpreted in a conventional manner given that the choice of window used to estimate the null-distribution parameters introduces an arbitrary scaling; thus, our choice of the *d*′ criterion to calculate CMR was instead based on avoiding floor and ceiling effects. Results indicate that the CMR circuit model shows a CMR effect consistent with actual cochlear nucleus data in that signal detectability is best in the Comodulated condition, followed by the Reference and Codeviant conditions (compare Figs. 4A and 4B with Figs. 2 and 6A from Pressnitzer et al., 2001, respectively). The size of the predicted CMR effect is also consistent with perceptual measurements (Mok et al., 2021). As expected, no CMR effect is seen at the level of the auditory nerve. Thus, the CMR circuit model accounts for the improved signal representation in the Comodulated condition where the masker is more easily segregable from the target signal, an advantage that derives from the fact that the different masking components are temporally coherent with one another. In addition, it also accounts for the greater cross-channel interference in the Codeviant condition, where the flanking components are temporally coherent with the target signal that is presented in the dips of the OFC. Finally, when the modulation rate of the input SAM tones was varied, CMR effects were still seen and followed the same low-pass trend as human perceptual data (Carlyon et al., 1989) (Fig. 4C).

**Figure 4:**
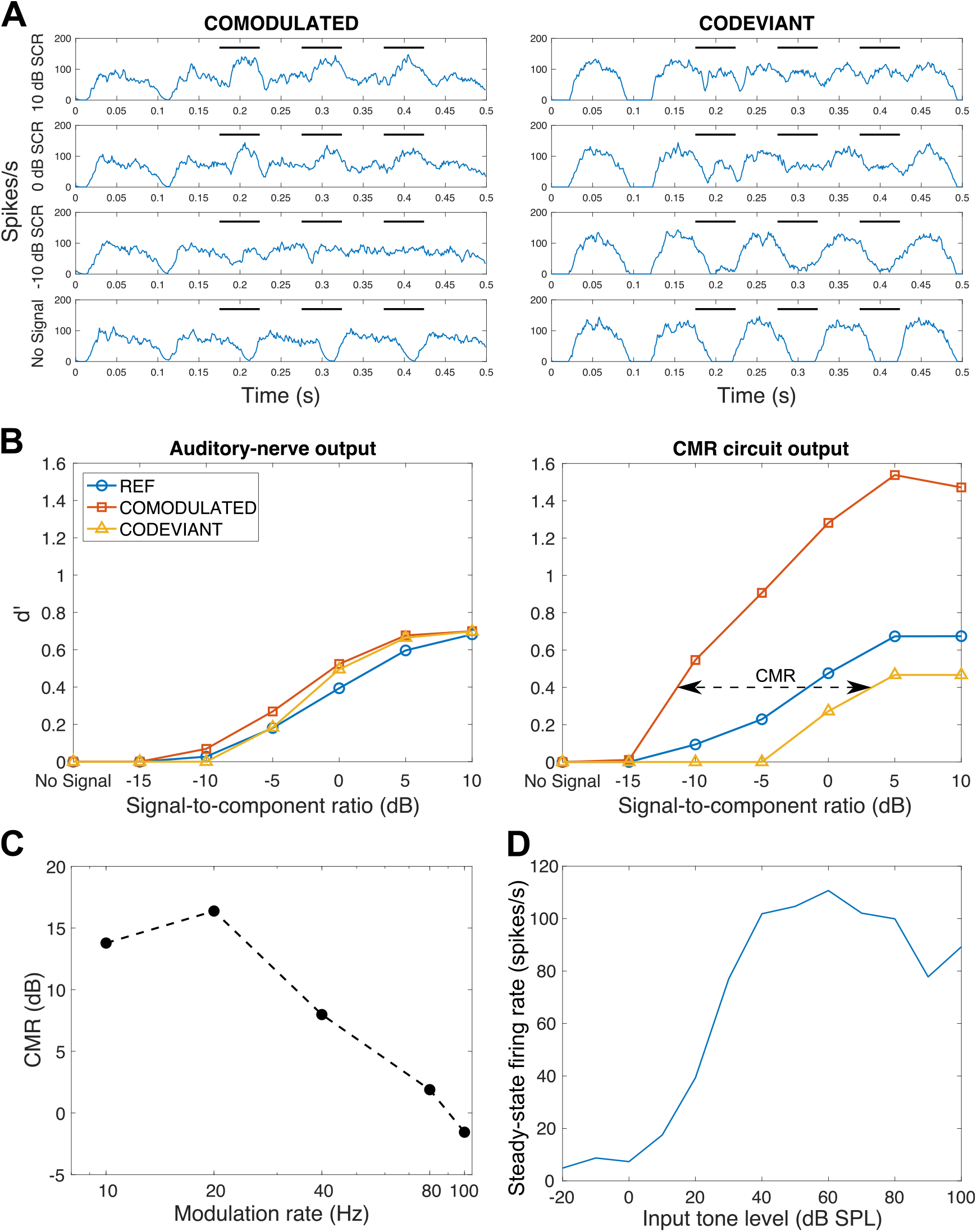
CMR circuit model validation. **Panel A** shows PSTH outputs from the CMR circuit model at 1.1 kHz CF for the stimuli in Figure 3. Results are shown separately for the Comodulated and Codeviant conditions, and at different SCRs. The black horizontal bars indicate the time points corresponding to when the target signal was presented. **Panel B** summarizes the results from Panel A by showing the neurometric sensitivity, *d*^*/*^, as a function of SCR for the auditory-nerve and CMR circuit model outputs (both at 1.1 kHz CF). The CMR circuit model shows a clear separation between the Comodulated and Codeviant conditions, i.e., a CMR effect. This is not seen at the level of the auditory nerve. **Panel C** shows the variation in the CMR obtained from the circuit model as a function of modulation rate. **Panel D** shows the pure-tone rate-level function (i.e., mean steady-state firing rate versus input tone level) for the CMR circuit model.

**Figure 5:**
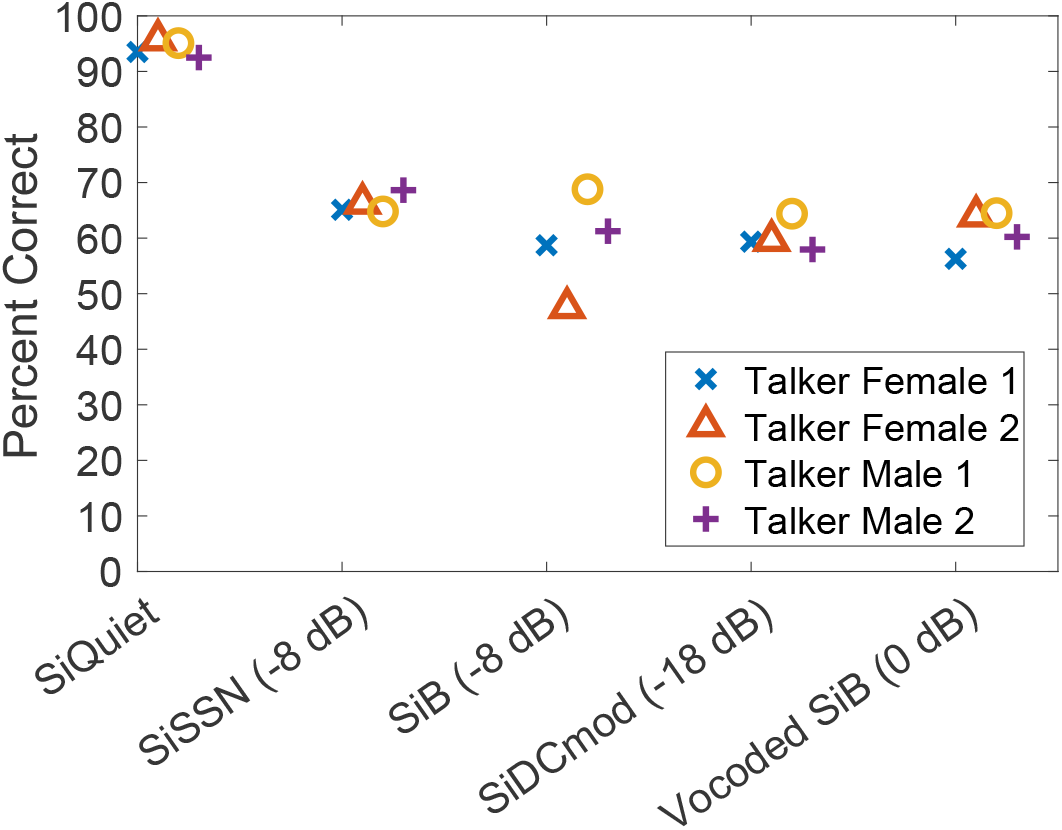
Overall intelligibility measured in the online consonant identification study for different conditions and talkers. Approximately equal overall intelligibility was achieved for SiSSN, SiDCmod, SiB, and Vocoded SiB (N=191).

**Figure 6:**
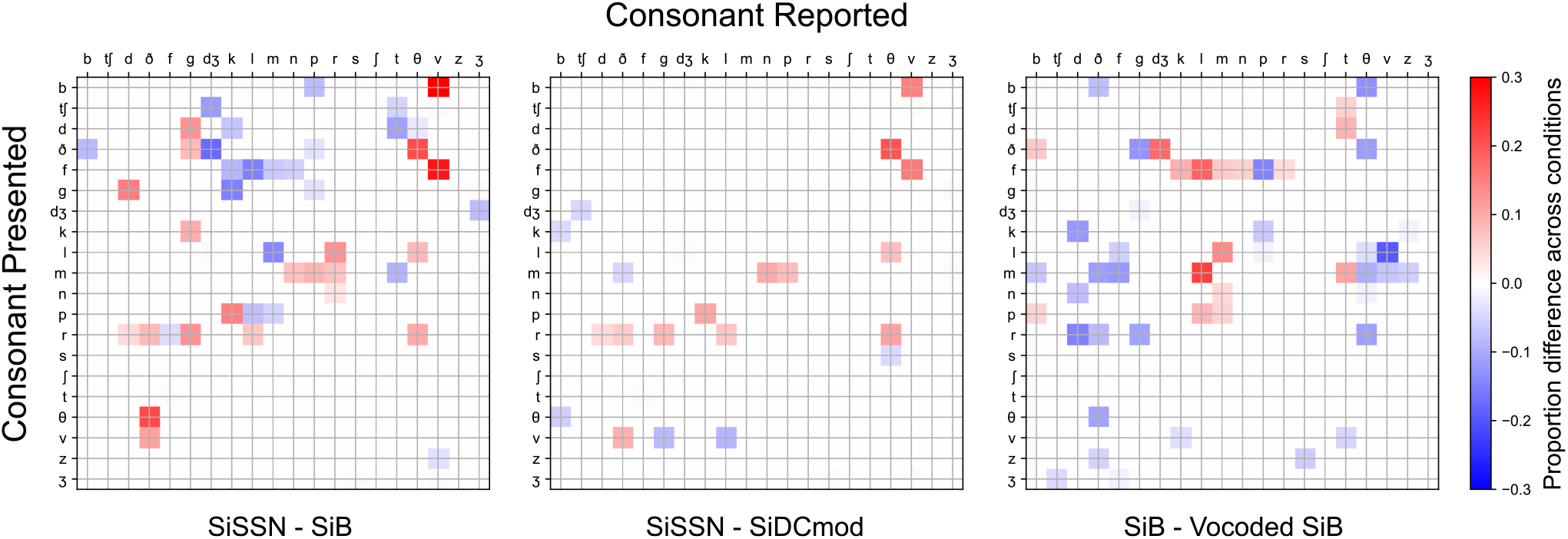
Measured consonant confusion-matrix differences across conditions (pooled over samples; N=191). The first two columns represent differences across maskers with different modulation spectra, whereas the third column shows the difference across stimuli with intact versus degraded TFS information. Only significant differences are shown, after permutation testing with multiple-comparisons correction (5% FDR). As the modulation statistics of the masker or the TFS content were varied, statistically significant differences emerged in the confusion patterns across conditions. Overall intelligibility was normalized to 60% for this analysis (Section 2.5) so that differences in confusion matrices across conditions could be attributed to changes in consonant categorization and category errors rather than differences in overall error counts (due to one condition being inherently easier at a particular SNR).

Each scene analysis model was separately *calibrated* by fitting a logistic/sigmoid function mapping the neural consonant confusion matrix entries from that model for the SiSSN condition to corresponding perceptual measurements. The mapping derived from this calibration was used to *predict* perceptual consonant confusion matrices from the corresponding neural confusion matrices for unseen conditions. Voicing, POA, and MOA confusion matrices were then derived by pooling over all consonants. Finally, the Pearson correlation coefficient was used to compare model predictions to perceptual measurements across the voicing, POA, and MOA categories. The prediction accuracy for the different models is reported in Section 3.

### 2.8 Statistical analysis

Permutation testing (Nichols and Holmes, 2002) with multiple-comparisons correction at 5% false-discovery rate (FDR; Benjamini and Hochberg, 1995) was used to extract significant differences in the SiSSN, SiB, SiDCmod, and Vocoded SiB consonant confusion matrices quantified in Section 2.5. The null distributions for permutation testing were obtained using a non-parametric shuffling procedure, which ensured that the data used in the computation of the null distributions had the same statistical properties as the measured confusion data. A separate null distribution was generated for each consonant. Each realization from each null distribution was obtained by following the same computations used to obtain the actual difference in the confusion matrices between conditions, but with random shuffling of condition labels corresponding to the measurements. This procedure was independently repeated with 10,000 distinct randomizations for each null distribution.

The p-values for the Pearson correlation coefficients between model predictions and perceptual measurements (Tables 3 and 4) were derived using Fisher’s approximation (Fisher, 1921).

**Table 3:** Pearson correlation coefficients between within-channel model predictions and perceptual measurements. Results are listed separately for the diagonal entries of the confusion matrix (i.e., proportion correct for the different consonant phonetic categories), off-diagonal entries (i.e., true confusions), and across all entries.

| Condition | Diagonal entries |  | Off-diagonal entries |  | All entries |  |
| --- | --- | --- | --- | --- | --- | --- |
|  | Correlation | p-value | Correlation | p-value | Correlation | p-value |
| SiB | 72% | 0.0026 ** | 64% | $10^{-7}$ *** | 87% | $10^{-21}$ *** |
| SiDCmod | 66% | 0.0072 ** | 64% | $10^{-7}$ *** | 83% | $10^{-17}$ *** |
| Vocoded SiB | 4% | 0.4445 | 40% | 0.0019 ** | 75% | $10^{-13}$ *** |
| SiSSN | 83% | 0.0002 *** | 67% | $10^{-8}$ *** | 87% | $10^{-21}$ *** |

**Table 4:** Pearson correlation coefficients between across-channel model predictions and perceptual measurements. Results are listed separately for the diagonal entries of the confusion matrix (i.e., proportion correct for the different consonant phonetic categories), off-diagonal entries (i.e., true confusions), and across all entries.

| Condition | Diagonal entries |  | Off-diagonal entries |  | All entries |  |
| --- | --- | --- | --- | --- | --- | --- |
|  | Correlation | p-value | Correlation | p-value | Correlation | p-value |
| SiB | 85% | 0.0001 *** | 73% | $10^{-10}$ *** | 90% | $10^{-24}$ *** |
| SiDCmod | 88% | $10^{-5}$ *** | 72% | $10^{-9}$ *** | 86% | $10^{-20}$ *** |
| Vocoded SiB | 63% | 0.0103 * | 70% | $10^{-9}$ *** | 86% | $10^{-20}$ *** |
| SiSSN | 89% | $10^{-5}$ *** | 81% | $10^{-13}$ *** | 92% | $10^{-27}$ *** |

To test whether the improvements in prediction accuracy (i.e., the correlation between model predictions and perceptual measurements) offered by the across-channel model compared to the within-channel model are statistically significant, a permutation procedure was employed once again. Under the null hypothesis that the within- and across-channel models are equivalent in their predictive power, the individual entries of the confusion matrices predicted by the two models can be swapped without effect on the results. Thus, to generate each realization of the null distribution of the correlation *improvement*, a randomly chosen half of the confusion matrix entries were swapped; this permutation procedure was independently repeated 100,000 times. A separate null distribution was generated in this manner for each condition. The actual improvements in correlation were compared against the corresponding null distributions to estimate (uncorrected) p-values. To adjust for multiple testing, an FDR procedure (Benjamini and Hochberg, 1995) was employed. Table 5 indicates whether each test met criteria for statistical significance under an FDR threshold of 5%.

**Table 5:**
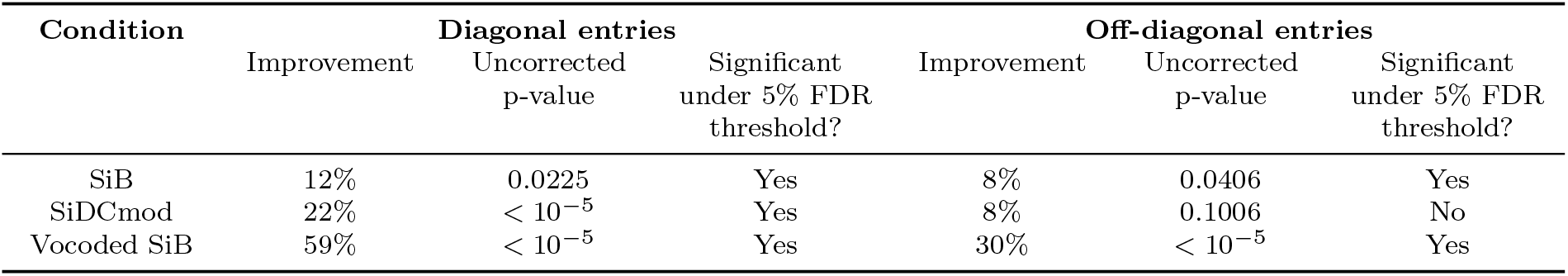
Improvement in prediction accuracy offered by the across-channel model compared to the within-channel model. The across-channel model showed improved correlations between model predictions and perceptual measurements for all of the unseen conditions, with the largest improvement apparent for Vocoded SiB.

### 2.9 Software accessibility

Subjects were directed from Prolific to the SNAPlab online psychoacoustics infrastructure (https://snaplabonline.com; Mok et al., 2021) to perform the study. Offline data analyses were performed using custom software in Python (Python Software Foundation, https://www.python.org) and MATLAB (The MathWorks, Inc., Natick, MA). Copies of all custom code can be obtained from the authors.

## 3 Results

Figure 5 shows speech intelligibility measurements from the online consonant identification study. Approximately equal overall intelligibility was achieved for SiSSN, SiDCmod, SiB, and Vocoded SiB due to our careful choice of SNRs for these conditions based on piloting (see Table 1). This was done to obtain roughly equal variance in the consonant confusion estimates for these conditions, which allows us to fairly compare confusion patterns across them. Equalizing intelligibility also maximizes the statistical power for detecting differences in the pattern of confusions. *∼*60% overall intelligibility was obtained in each condition, which yielded a sufficient number of confusions for analysis.

Given that all psychophysical data were collected online, data quality was verified by comparing results for SiSSN with previous lab-based findings; the analyses performed and the results are described in Mok et al. (2021) and Viswanathan et al. (2021b).

The identification data collected in the Test stage of the online experiment were used to construct a consonant confusion matrix for each condition (see Section 2.5). Then, statistically significant differences in these matrices across conditions were extracted (Section 2.8). Results (Fig. 6) show significant differences in the confusion patterns across (i) conditions with different masker modulation statistics, and (ii) stimuli with intact versus degraded TFS information. Computational modeling was then used to predict these differences across conditions to test specific theories of scene analysis (Section 2.7).

Figure 7 shows results from the calibration step of testing the within- and across-channel models of scene analysis. In this step, a separate logistic/sigmoid function was fit for each model to map neural confusion matrix entries for the SiSSN condition to corresponding perceptual measurements.

**Figure 7:**
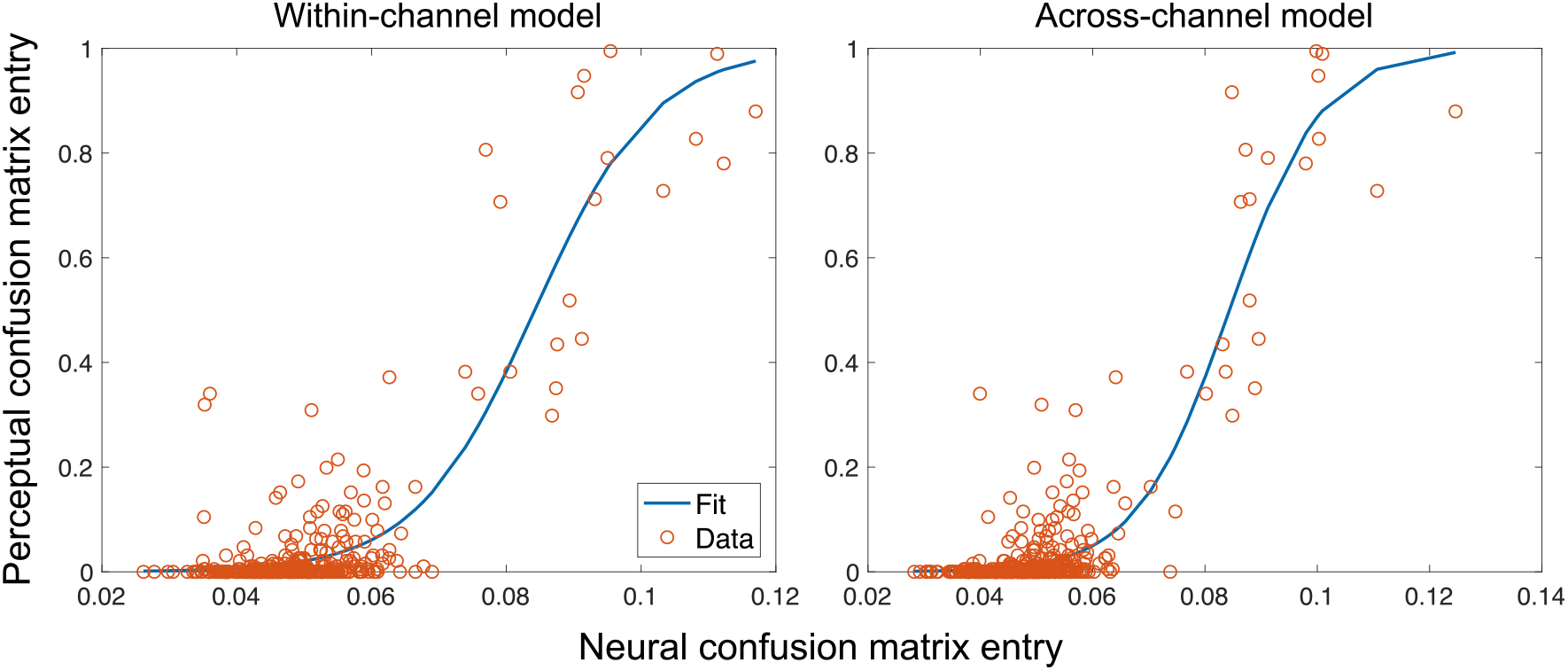
Calibration result for the within- and across-channel models of scene analysis. Shown are the model-specific sigmoid/logistic functions that were fit to map neural confusion matrix entries for the SiSSN condition to corresponding perceptual measurements. This model-specific mapping was used to predict perceptual consonant confusion matrices from neural confusion matrices for unseen conditions.

The model-specific mapping derived in the calibration step was used to predict perceptual consonant confusion matrices for each of the scene analysis models from the neural confusion matrices for unseen conditions (not used in calibration). Then, voicing, POA, and MOA confusion matrices were derived by pooling over all consonants (Figs. S1, S2, and S3). Finally, model predictions were compared to perceptual measurements for the different confusion matrix entries across the voicing, POA, and MOA categories. The results are shown in Figure 8 for SiB, SiDCmod, and Vocoded SiB. The SiQuiet condition is not visualized, as there were ceiling effects in the intelligibility measurements (i.e., the diagonal entries of the confusion matrix were dominant) and very few confusions (i.e., off-diagonal entries were rare), which made it infeasible to meaningfully evaluate the quality of predictions for this condition (as there was no variance across either the on-or off-diagonals). But overall, across all entries for SiQuiet, both models predicted diagonal entries close to one and off-diagonal entries close to zero, in line with perceptual measurements.

**Figure 8:**
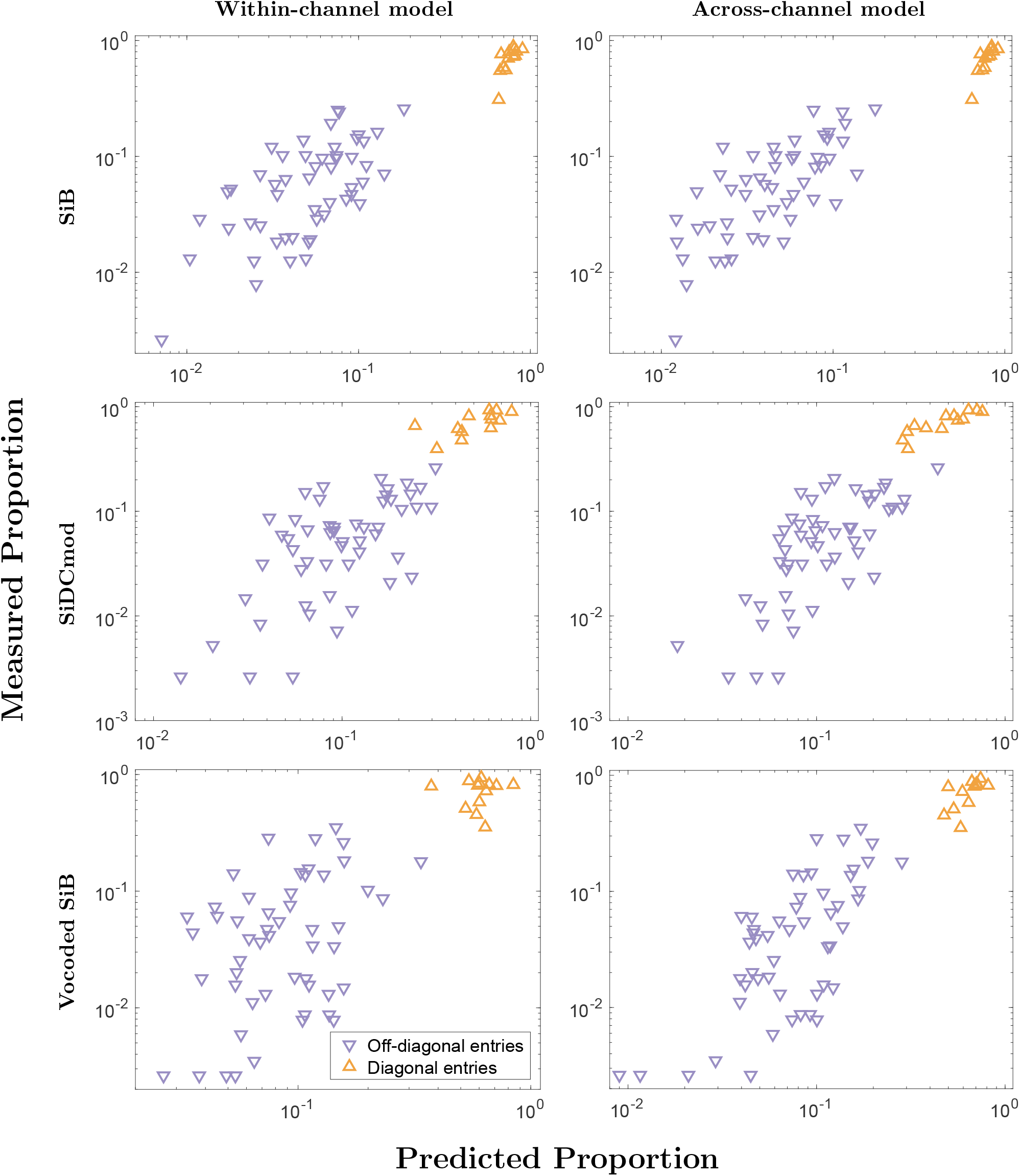
Within- and across-channel model predictions versus measured confusion matrix entries for the unseen conditions. Diagonal entries correspond to intelligibility measurements for the different consonant phonetic categories (transmission scores for voicing, POA, and MOA), and off-diagonal entries correspond to true confusions. It can be seen that the cluster of points is less dispersed for the across-channel model compared to the within-channel model, indicating greater predictive accuracy. These trends are quantified in Tables 3, 4, and 5.

Pearson correlation coefficients were computed between the model predictions and perceptual measurements (shown in Fig. 8) and are given in Tables 3 and 4 for the within- and across-channel models, respectively. Since the range of confusion matrix entries spanned three orders of magnitude, all comparisons were performed with log-transformed values. The correlations were statistically significant across all non-vocoded conditions for the within-channel model, and across all conditions for the across-channel model (see Section 2.8 for statistical analysis details). The strong correlation of the within-channel model predictions with perceptual data in the non-vocoded conditions (where TFS cues are preserved) provides independent evidence that speech understanding is strongly influenced by modulation masking when TFS cues are available (Viswanathan et al., 2021a); moreover, this result also suggests that modulations are used differently by the brain in the absence of natural TFS.

The across-channel model produced stronger correlation values compared to the within-channel model for all conditions, and the improvements were statistically significant across all conditions even after correcting for multiple comparisons (Table 5; for analysis details, see Section 2.8). Thus, a simple physiologically plausible model of across-channel cochlear nucleus processing that shows CMR (Fig. 4) also yields category confusion predictions that match behavioral data, and more specifically improves predictions compared to a within-channel model. Note that our within-channel model assumes perfect segregability of target-masker components that are separated in CF and MF (in line with current speech-intelligibility models; Jørgensen et al., 2013; Relaño-Iborra et al., 2016), and only models within-channel modulation masking. Specifically, within a particular channel (i.e., CF) and MF, masker modulations that are not in phase with the target are the only components that mask the target. However, our across-channel model simulates both within-channel modulation masking and cross-channel temporal-coherence-based interference. Specifically, masker components that are in a different channel from the target but that are temporally coherent with the target can interfere with target coding and perception. We implemented this interference via the CMR circuit model (Fig. 1) where temporally coherent pieces of the target and masker, even across distinct cochlear channels, coherently drive the wideband inhibitor (WBI), thereby enhancing outputs of the narrowband (NB) unit (which is inhibited by the WBI) that are incoherent with the masker. Thus, our finding that model predictions are improved when cross-channel processing is added is consistent with the theory that across-channel temporal coherence shapes scene analysis (Elhilali et al., 2009). Moreover, this result also suggests that physiological computations that exist as early as the cochlear nucleus can contribute significantly to temporal-coherence-based scene analysis. Note that improvements to confusion predictions are apparent with the across-channel model for the same range of model parameters for which the CMR effect is also apparent.

Another key result from Table 5 is that the condition that showed the greatest improvement in confusion matrix predictions between the within- and across-channel models is Vocoded SiB. The masker in Vocoded SiB produces both within-channel modulation masking and cross-channel interference (as described above). These masking and interference effects are partially mitigated in intact SiB (and other non-vocoded conditions) compared to Vocoded SiB, because the brain can use the pitch cue supplied by natural TFS to better separate the target and masker (Darwin, 1997; Oxenham and Simonson, 2009). The across-channel model is a better fit to perceptual data for all conditions, which suggests that cross-channel interference affects perceptual data. Thus, the improvement offered by this model is likely most apparent for vocoded SiB because cross-channel interference effects contribute most to perception in this condition.

Note that while the main difference between the two scene analysis models tested in the current study is the exclusion/inclusion of cross-channel processing, another difference is that the within-channel model discards TFS, whereas the across-channel model uses the full simulated auditory-nerve output to drive the CMR circuit model. This raises the possibility that part of the improvement offered by the across-channel model could come simply from the inclusion of TFS information within each channel independently. To investigate whether the poorer performance of the within-channel model was partly due to discarding TFS, we re-ran the within-channel model by retaining the full auditory-nerve output (results not shown). We found that the predictions from the modified within-channel model were not significantly better than the original within-channel model. This confirms that the improvement in predictions given by the across-channel model comes largely from across-channel CMR effects, suggesting that categorical perception is sensitive to the temporal coherence across channels. Moreover, these CMR effects were restricted to low rates (*<* 80 Hz or so; Fig. 4C), consistent with perceptual data (Carlyon et al., 1989). This suggests that the cross-channel processing did not benefit much from the TFS information included in driving the CMR circuit model.

## 4 Discussion

To probe the contribution of temporal-coherence processing to speech understanding in noise, the present study used a behavioral experiment to measure consonant identification in different masking conditions in conjunction with physiologically plausible computational modeling. To the best of our knowledge, this is the first study to use confusion patterns in speech categorization to test theories of auditory scene analysis. The use of confusion data provides independent constraints on our understanding of scene-analysis mechanisms beyond what overall intelligibility can provide. This is because percent correct data only convey binary information about whether or not target coding was intact, whereas consonant categorization and confusion data provide richer information about what sound elements received perceptual weighting.

We constructed computational models simulating (i) purely within-channel modulation masking (in line with current speech-intelligibility models; Relaño-Iborra et al., 2016), and (ii) a combination of within-channel modulation masking and across-channel temporal-coherence processing mirroring physiological computations that are known to exist in the cochlear nucleus (Pressnitzer et al., 2001). Our across-channel temporal coherence circuit produced a CMR effect (Fig. 4) that is consistent with actual cochlear nucleus data (Pressnitzer et al., 2001) and perceptual measurements (Mok et al., 2021). Moreover, consonant confusion pattern predictions were significantly improved for all tested conditions with the addition of this cross-channel processing (Table 5), which suggests that temporal-coherence processing strongly shapes speech categorization when listening in noise. This result is consistent with the theory that comodulated features of a sound source are perceptually grouped together, and that masker elements that are temporally coherent with target speech but in a different channel from the target perceptually interfere (Darwin, 1997; Schooneveldt and Moore, 1987; Apoux and Bacon, 2008). The only case where the within- and across-channel models were statistically equivalent was in predicting the off-diagonal entries (i.e., true confusions) for the SiDCmod condition; this may be because this condition has little coherent cross-channel interference from the masker as the masker is unmodulated (Stone et al., 2012).

An important difference between the cross- and within-channel masking simulated in our models is that while the cross-channel interference was produced by masker fluctuations that were temporally coherent with the target, the within-channel masking was produced by masker components that were matched in both CF and MF with target components. While current speech-intelligibility models simulate the latter type of masking (Jørgensen et al., 2013; Relaño-Iborra et al., 2016), they do not account for cross-channel temporal-coherence-based masking as we have done here. This may explain why these models fail in certain conditions, including for vocoded stimuli (Steinmetzger et al., 2019). Indeed, even in the present study, although our within-channel modulation masking model reasonably accounted for category confusions, it failed when TFS cues were unavailable (Table 3). One explanation for this is that because pitch-based masking release is poorer in the vocoded condition due to degraded TFS information (Oxenham and Simonson, 2009), the effects of cross-channel interference are more salient. This may also be the reason why the Vocoded SiB condition showed the greatest improvement in confusion pattern predictions after adding cross-channel processing (Table 5), which models these interference effects.

Although the lateral inhibition network used in Elhilali et al. (2003) bears some similarities to the across-channel CMR circuit model used in the current study, the CMR circuit model was explicitly based on physiological computations present in the cochlear nucleus and their CMR properties. Thus, another implication of the results of the present study is that physiological computations that exist as early as the cochlear nucleus can contribute significantly to temporal-coherence-based scene analysis. Such effects likely accumulate as we ascend along the auditory pathway (Elhilali et al., 2009; Teki et al., 2013; O’Sullivan et al., 2015). Note that the CMR circuit model does not perform pitch-range temporal-coherence processing and no CMR effect was seen at high modulation rates (Fig. 4C), consistent with perceptual data in the literature (Carlyon et al., 1989). Despite this, our across-channel model significantly improved predictions of category confusions compared to the within-channel model, which suggests that temporal-coherence processing at lower modulation rates is perceptually important. A future research direction is to extend the modeling framework proposed here to study the contributions of scene-analysis mechanisms beyond the specific aspects of temporal-coherence processing studied here. One such extension could be to account for pitch-based source segregation (Bregman, 1990), perhaps by modeling a combined temporal-place code for pitch processing (Oxenham et al., 2004; Shamma and Klein, 2000; Oxenham and Simonson, 2009).

One limitation of the periphery model we used (Bruce et al., 2018) is that it was developed to match nerve responses to simple stimuli. However, this family of periphery models has been successfully used to account for complex phenomena such as synchrony capture (Delgutte and Kiang, 1984), formant coding in the midbrain (Carney et al., 2015), and qualitative aspects of evoked potentials such as auditory brainstem responses and frequency-following responses (Shinn-Cunningham et al., 2013). Although a debate exists regarding the spatio-temporal properties of different periphery models in cochlear responses (Verhulst et al., 2015; Vecchi et al., 2021), those differences are subtle compared to the slower CMR effects that are important for the present study. A more general limitation of the models used in this study is that they are simple and do not incorporate many aspects of speech perception (e.g., context effects; Dubno and Levitt, 1981) because the goal here is to test specific theories of scene analysis. Nevertheless, the contrast between the models would be unaffected by these higher-order effects.

## 5 Acknowledgments

This research was supported by grants from the National Institutes of Health [F31DC017381 (to V.V.) and R01DC009838 (to M.G.H.)] and Office of Naval Research [ONR N00014-20-12709 (to B.G.S.-C.)]. We thank Hari Bharadwaj for access to online psychoacoustics infrastructure (https://snaplabonline.com; Mok et al., 2021). We also thank Andrew Sivaprakasam, François Deloche, Hari Bharadwaj, and Ravinderjit Singh for valuable feedback on an earlier version of this manuscript.

## 6 Author Contributions

V.V., B.G.S.-C., and M.G.H. designed research; V.V. performed research; V.V. analyzed data; V.V. wrote the paper with edits from B.G.S.-C. and M.G.H.

## 7 Supplementary Information

For completeness, the full set of model-predicted and measured perceptual confusion matrices are shown in Figures S1, S2, and S3 for the voicing, POA, and MOA categories, respectively. Results are shown only for the SiB, SiDCmod, and Vocoded SiB conditions (i.e., the conditions unseen by the calibration step and having a sufficient number of confusions for prediction).

---

**Figure S1:**
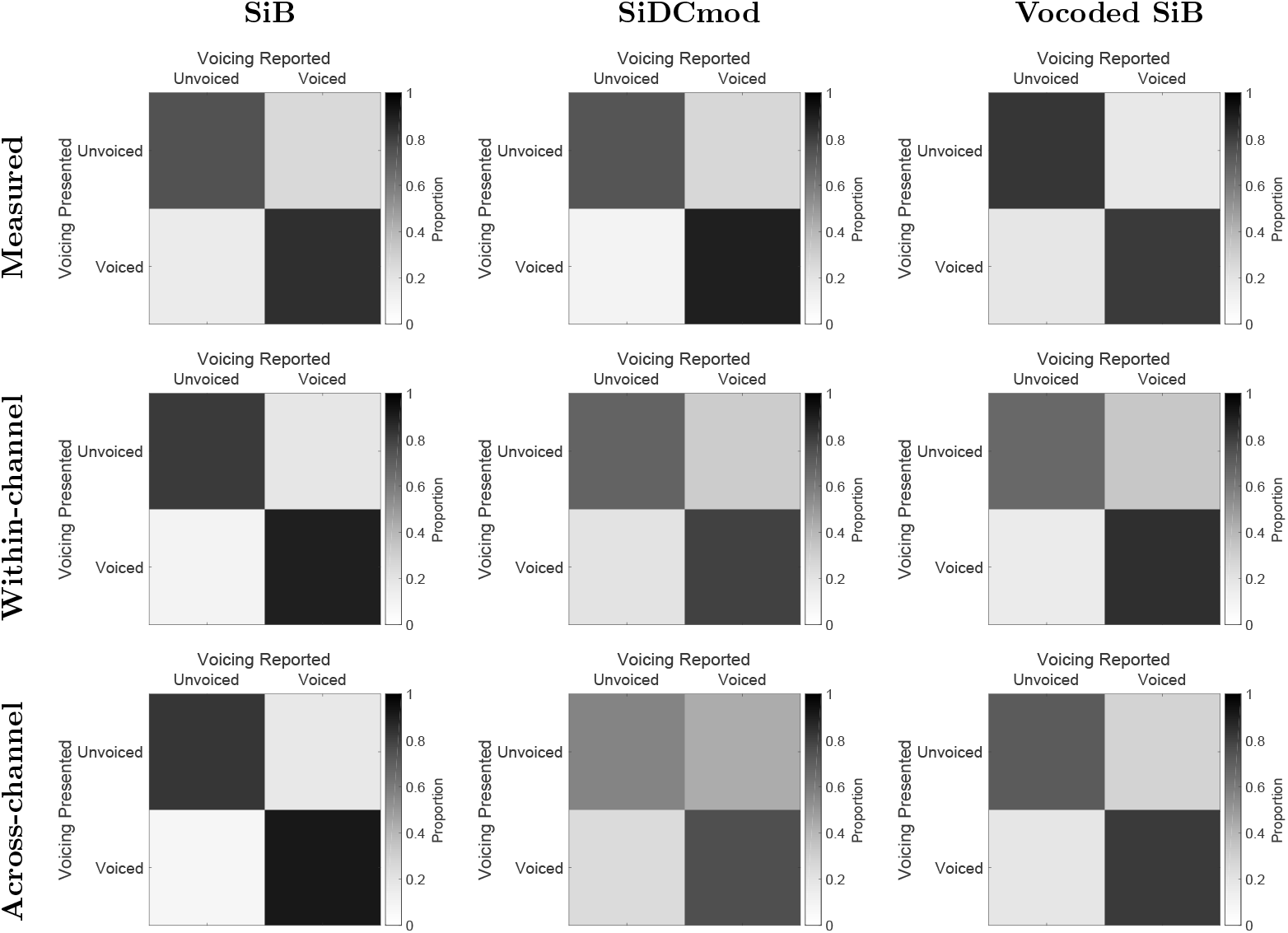
Full set of measured (top row) and model-predicted (middle and bottom rows) voicing confusion matrices.

**Figure S2:**
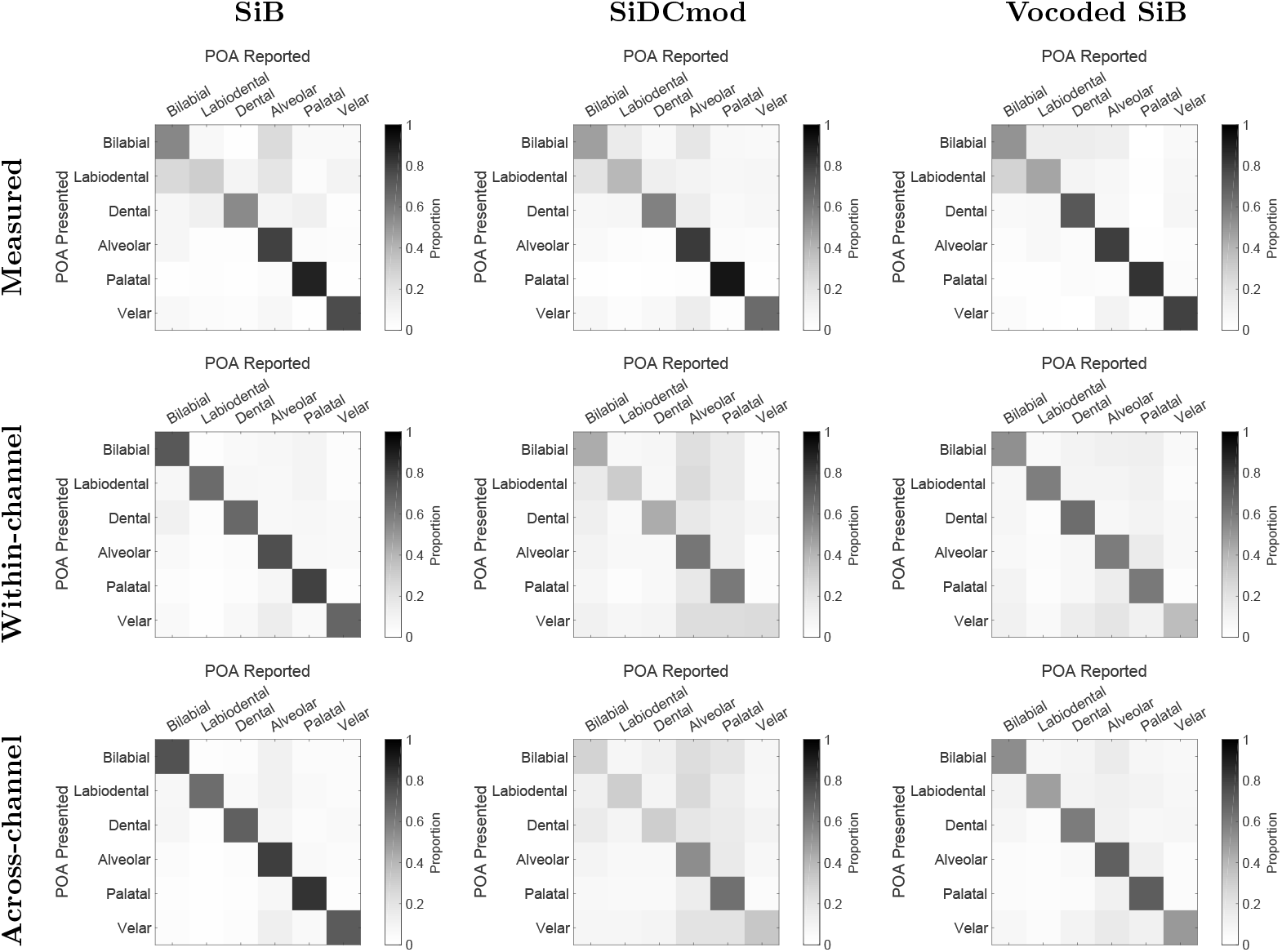
Full set of measured (top row) and model-predicted (middle and bottom rows) place of articulation (POA) confusion matrices.

**Figure S3:**
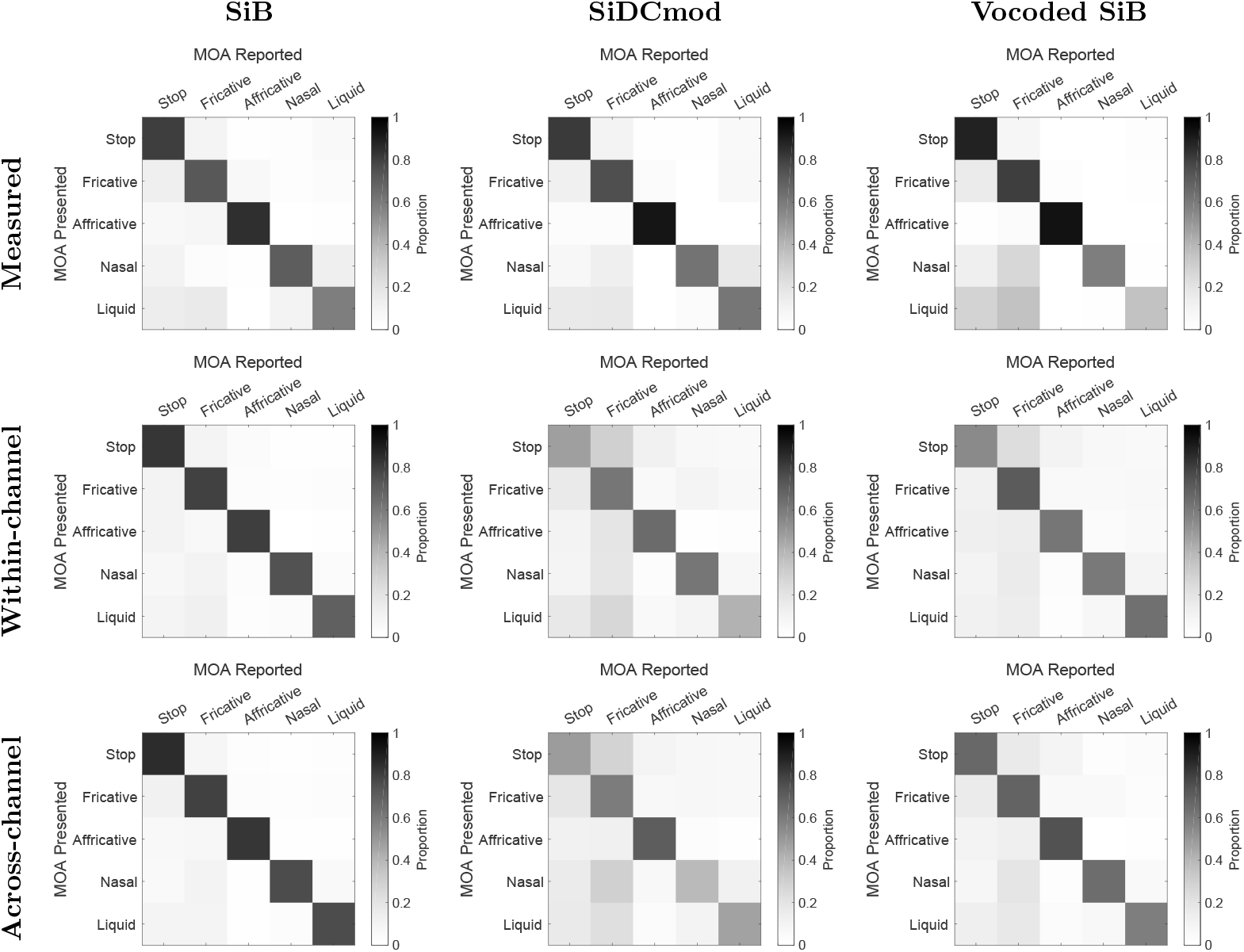
Full set of measured (top row) and model-predicted (middle and bottom rows) manner of articulation (MOA) confusion matrices.

## References

Apoux, F. and Bacon, S. P. (2008). Selectivity of modulation interference for consonant identification in normal-hearing listeners. J Acoust Soc Am, 123(3):1665–1672.

Bacon, S. P. and Grantham, D. W. (1989). Modulation masking: Effects of modulation frequency, depth, and phase. J Acoust Soc Am, 85(6):2575–2580.

Benjamini, Y. and Hochberg, Y. (1995). Controlling the false discovery rate: a practical and powerful approach to multiple testing. J Royal Stat Soc Series B Stat Methodol, pages 289–300.

Bregman, A. (1990). Auditory scene analysis: the perceptual organization of sound. Cambridge, MA: MIT Press.

Bruce, I. C., Erfani, Y., and Zilany, M. S. (2018). A phenomenological model of the synapse between the inner hair cell and auditory nerve: Implications of limited neurotransmitter release sites. Hear Res, 360:40–54.

Carlyon, R. P., Buus, S., and Florentine, M. (1989). Comodulation masking release for three types of modulator as a function of modulation rate. Hear Res, 42(1):37–45.

Carney, L. H., Li, T., and McDonough, J. M. (2015). Speech coding in the brain: representation of vowel formants by midbrain neurons tuned to sound fluctuations. Eneuro, 2(4).

Crouzet, O. and Ainsworth, W. A. (2001). On the various influences of envelope information on the perception of speech in adverse conditions: An analysis of between-channel envelope correlation. In Workshop on Consistent and Reliable Acoustic Cues for Sound Analysis, Aalborg, Denmark.

Darwin, C. J. (1997). Auditory grouping. Trends Cogn Sci, 1(9):327–333.

Delgutte, B. and Kiang, N. Y. (1984). Speech coding in the auditory nerve: I. vowel-like sounds. J Acoust Soc Am, 75(3):866–878.

Dubno, J. R. and Levitt, H. (1981). Predicting consonant confusions from acoustic analysis. J Acoust Soc Am, 69(1):249–261.

Elhilali, M., Chi, T., and Shamma, S. A. (2003). A spectro-temporal modulation index (stmi) for assessment of speech intelligibility. Speech Commun, 41(2-3):331–348.

Elhilali, M., Ma, L., Micheyl, C., Oxenham, A. J., and Shamma, S. A. (2009). Temporal coherence in the perceptual organization and cortical representation of auditory scenes. Neuron, 61(2):317–329.

Ewert, S. D. and Dau, T. (2000). Characterizing frequency selectivity for envelope fluctuations. J Acoust Soc Am, 108(3):1181–1196.

Fisher, R. A. (1921). On the ‘probable error’ of a coefficient of correlation deduced from a small sample. Metron, 1:1–32.

Glasberg, B. R. and Moore, B. C. (1990). Derivation of auditory filter shapes from notched-noise data. Hear Res, 47(1):103–138.

Heinz, M. G., Colburn, H. S., and Carney, L. H. (2002). Quantifying the implications of nonlinear cochlear tuning for auditory-filter estimates. J Acoust Soc Am, 111(2):996–1011.

Hilbert, D. (1906). Grundzüge einer allgemeinen Theorie der linearen Integralgleichungen. Vierte Mitteilung. Nachrichten von der Gesellschaft der Wissenschaften zu Göttingen, Mathematisch-Physikalische Klasse, 1906:157–228.

Hillyard, S. A., Hink, R. F., Schwent, V. L., and Picton, T. W. (1973). Electrical signs of selective attention in the human brain. Science, 182(4108):177–180.

Jørgensen, S., Ewert, S. D., and Dau, T. (2013). A multi-resolution envelope-power based model for speech intelligibility. J Acoust Soc Am, 134(1):436–446.

Killion, M. C., Niquette, P. A., Gudmundsen, G. I., Revit, L. J., and Banerjee, S. (2004). Development of a quick speech-in-noise test for measuring signal-to-noise ratio loss in normal-hearing and hearing-impaired listeners. J Acoust Soc Am, 116(4):2395–2405.

Krishnan, L., Elhilali, M., and Shamma, S. (2014). Segregating complex sound sources through temporal coherence. PLoS Comput Biol, 10(12):e1003985.

Meddis, R., Delahaye, R., O’Mard, L., Sumner, C., Fantini, D. A., Winter, I., and Pressnitzer, D. (2002). A model of signal processing in the cochlear nucleus: comodulation masking release. Acta Acust united Ac, 88(3):387–398.

Miller, G. A. and Nicely, P. E. (1955). An analysis of perceptual confusions among some english consonants. J Acoust Soc Am, 27(2):338–352.

Mok, B. A., Viswanathan, V., Borjigin, A., Singh, R., Kafi, H. I., and Bharadwaj, H. M. (2021). Web-based psychoacoustics: Hearing screening, infrastructure, and validation. bioRxiv, DOI: 10.1101/2021.05.10.443520.

Nichols, T. E. and Holmes, A. P. (2002). Nonparametric permutation tests for functional neuroimaging: a primer with examples. Hum Brain Mapp, 15(1):1–25.

O’Sullivan, J. A., Shamma, S. A., and Lalor, E. C. (2015). Evidence for neural computations of temporal coherence in an auditory scene and their enhancement during active listening. J Neurosci, 35(18):7256–7263.

Oxenham, A. J., Bernstein, J. G., and Penagos, H. (2004). Correct tonotopic representation is necessary for complex pitch perception. Proc Natl Acad Sci USA, 101(5):1421–1425.

Oxenham, A. J. and Shera, C. A. (2003). Estimates of human cochlear tuning at low levels using forward and simultaneous masking. J Assoc Res Otolaryngol, 4(4):541–554.

Oxenham, A. J. and Simonson, A. M. (2009). Masking release for low-and high-pass-filtered speech in the presence of noise and single-talker interference. J Acoust Soc Am, 125(1):457–468.

Patterson, R. D., Nimmo-Smith, I., Holdsworth, J., and Rice, P. (1987). An efficient auditory filterbank based on the gammatone function. In a meeting of the IOC Speech Group on Auditory Modelling at RSRE, volume 2.

Pressnitzer, D., Meddis, R., Delahaye, R., and Winter, I. M. (2001). Physiological correlates of co-modulation masking release in the mammalian ventral cochlear nucleus. J Neurosci, 21(16):6377–6386.

Rabiner, L. (1993). Fundamentals of speech recognition. Fundamentals of speech recognition.

Relaño-Iborra, H., May, T., Zaar, J., Scheidiger, C., and Dau, T. (2016). Predicting speech intelligibility based on a correlation metric in the envelope power spectrum domain. J Acoust Soc Am, 140(4):2670–2679.

Schooneveldt, G. P. and Moore, B. C. (1987). Comodulation masking release (cmr): Effects of signal frequency, flanking-band frequency, masker bandwidth, flanking-band level, and monotic versus dichotic presentation of the flanking band. J Acoust Soc Am, 82(6):1944–1956.

Shamma, S. and Klein, D. (2000). The case of the missing pitch templates: how harmonic templates emerge in the early auditory system. J Acoust Soc Am, 107(5):2631–2644.

Shera, C. A., Guinan, J. J., and Oxenham, A. J. (2002). Revised estimates of human cochlear tuning from otoacoustic and behavioral measurements. Proc Natl Acad Sci U S A, 99(5):3318–3323.

Shinn-Cunningham, B. (2008). Object-based auditory and visual attention. Trends Cogn Sci, 12(5):182–186.

Shinn-Cunningham, B., Ruggles, D. R., and Bharadwaj, H. (2013). How early aging and environment interact in everyday listening: from brainstem to behavior through modeling. In Basic aspects of hearing, pages 501–510. Springer.

Singer, W. and Gray, C. M. (1995). Visual feature integration and the temporal correlation hypothesis. Annu Rev Neurosci, 18(1):555–586.

Steinmetzger, K., Zaar, J., Relaño-Iborra, H., Rosen, S., and Dau, T. (2019). Predicting the effects of periodicity on the intelligibility of masked speech: An evaluation of different modelling approaches and their limitations. J Acoust Soc Am, 146(4):2562–2576.

Stone, M. A., Füllgrabe, C., and Moore, B. C. (2012). Notionally steady background noise acts primarily as a modulation masker of speech. J Acoust Soc Am, 132(1):317–326.

Stone, M. A. and Moore, B. C. (2014). On the near non-existence of “pure” energetic masking release for speech. J Acoust Soc Am, 135(4):1967–1977.

Swaminathan, J. and Heinz, M. G. (2011). Predicted effects of sensorineural hearing loss on across-fiber envelope coding in the auditory nerve. J Acoust Soc Am, 129(6):4001–4013.

Teki, S., Chait, M., Kumar, S., Shamma, S., and Griffiths, T. D. (2013). Segregation of complex acoustic scenes based on temporal coherence. Elife, 2:e00699.

Vecchi, A. O., Varnet, L., Carney, L. H., Dau, T., Bruce, I. C., Verhulst, S., and Majdak, P. (2021). A comparative study of eight human auditory models of monaural processing.

Verhulst, S., Bharadwaj, H. M., Mehraei, G., Shera, C. A., and Shinn-Cunningham, B. G. (2015). Functional modeling of the human auditory brainstem response to broadband stimulation. J Acoust Soc Am, 138(3):1637–1659.

Viswanathan, V., Bharadwaj, H. M., Shinn-Cunningham, B. G., and Heinz, M. G. (2021a). Modulation masking and fine structure shape neural envelope coding to predict speech intelligibility across diverse listening conditions. bioRxiv, DOI: 10.1101/2021.03.26.437273.

Viswanathan, V., Shinn-Cunningham, B. G., and Heinz, M. G. (2021b). Temporal fine structure influences voicing confusions for consonant identification in multi-talker babble. bioRxiv, DOI: 10.1101/2021.05.11.443678.

Winter, I. M. and Palmer, A. R. (1990). Responses of single units in the anteroventral cochlear nucleus of the guinea pig. Hear Res, 44(2-3):161–178.

Zurek, P. M. (1993). Binaural advantages and directional effects in speech intelligibility. Acoustical factors affecting hearing aid performance, 2:255–275.

